# A biologically oriented algorithm for spatial sound segregation

**DOI:** 10.1101/2020.11.04.368548

**Authors:** Kenny F Chou, Virginia Best, H Steven Colburn, Kamal Sen

**Affiliations:** Department of Biomedical Engineering, Boston University, Boston, Massachusetts, United States of America; Department of Speech, Language and Hearing Sciences, Boston University, Boston, Massachusetts, United States of America

## Abstract

Listening in an acoustically cluttered scene remains a difficult task for both machines and hearing-impaired listeners. Normal-hearing listeners accomplish this task with relative ease by segregating the scene into its constituent sound sources, then selecting and attending to a target source. An assistive listening device that mimics the biological mechanisms underlying this behavior may provide an effective solution for those with difficulty listening in acoustically cluttered environments (e.g., a cocktail party). Here, we present a binaural sound segregation algorithm based on a hierarchical network model of the auditory system. In the algorithm, binaural sound inputs first drive populations of neurons tuned to specific spatial locations and frequencies. Lateral inhibition then sharpens the spatial response of the neurons. Finally, the spiking response of neurons in the output layer are then reconstructed into audible waveforms via a novel reconstruction method. We evaluate the performance of the algorithm with psychoacoustic measures of normal-hearing listeners. This two-microphone algorithm is shown to provide listeners with perceptual benefit similar to that of a 16-microphone acoustic beamformer in a difficult listening task. Unlike deep-learning approaches, the proposed algorithm is biologically interpretable and does not need to be trained on large datasets. This study presents a biologically based algorithm for sound source segregation as well as a method to reconstruct highly intelligible audio signals from spiking models.

**Author Summary:** Animal and humans can navigate complex auditory environments with relative ease, attending to certain sounds while suppressing others. Normally, various sounds originate from various spatial locations. This paper presents an algorithmic model to perform sound segregation based on how animals make use of this spatial information at various stages of the auditory pathway. We showed that the performance of this two-microphone algorithm provides as much benefit to normal-hearing listeners a multi-microphone algorithm. Unlike mathematical and machine-learning approaches, our model is fully interpretable and does not require training with large datasets. Such an approach may benefit the design of machine hearing algorithms. To interpret the spike-trains generated in the model, we designed a method to recover sounds from model spikes with high intelligibility. This method can be applied to spiking neural networks for audio-related applications, or to interpret each node within a spiking model of the auditory cortex.

## Introduction

Attending to a single conversation partner in the presence of multiple distracting talkers (i.e., the Cocktail Party Problem, CPP) is a complicated and difficult task for machines and humans [1–3]. While some normal-hearing listeners can accomplish this task with relative ease, other groups of listeners report great difficulty – such as those with sensorineural hearing loss [4–6], cochlear implant users [7–10], subgroups of children [11] and adults with “hidden hearing loss” [12–14]. At a cocktail party, talkers are distributed in space, and normal-hearing listeners appear to make use of spatial cues (i.e., interaural timing and level differences, or ITDs and ILDs, respectively) to perceptually localize and segregate sound mixtures into individual spatial components. Indeed, spatial listening has been shown to provide enormous benefit to listeners in cocktail-party scenarios [15,16].

Spatial processing also plays a key role in several potential solutions to the CPP. For example, acoustic beamforming techniques utilize multiple microphones to selectively enhance signals from a desired direction [17,18], and is often employed in hearing aids [19–23]. Some “super-directional” beamforming algorithms, which employ microphone arrays [24–26], have been shown to provide large benefits to listeners with hearing impairments [20,27], especially when combined with spatial-cue preservation strategies [28,29].

Unlike beamformers, typical computational auditory scene analysis (CASA) algorithms use only one or two microphones to perform sound segregation. This class of algorithms decompose sound streams into segments of small time and frequency chunks, then group together segments that may originate from the same sound source based on biologically relevant features, such as ITDs and ILDs [30]. Research in the CASA domain has moved toward using deep neural networks (DNNs) to perform grouping due to their high performance over other classes of grouping algorithms [3,31,32]; however, the performance of DNNs depends on supervised training on large sets of hand-labeled data and may not generalize to test examples outside the training set [32,33], thus limiting their practicality for hearing-aid processing.

The normal-functioning auditory system has several advantages compared to both DNNs and super-directional beamformers because it does not require *a priori* knowledge of the location or number of sources, it generalizes to any environment, and it only requires two input channels – the left and right ears. We recently proposed a physiologically based algorithm (PA) for sound segregation. The PA mimicked the normal auditory system by constructing model neurons that are sensitive to specific locations in space [34]. Although the PA provided proof-of-concept for a biologically based CASA algorithm, its performance was limited by audio reconstructions with low quality and intelligibility, and it did not preserve binaural cues [34]. These drawbacks limited the practical use of the PA for applications in hearing-assistive devices and machine hearing.

In this study, we present a biologically-oriented sound segregation algorithm (BOSSA) with the ultimate goal of helping listeners converse in multi-talker environments by spatially segregating the embedded talkers and isolating the talker of interest. BOSSA extends the PA by improving the spatial response of model neurons and by introducing a novel reconstruction method to convert neural spike ensembles back into intelligible audio waveforms. We compared the proposed two-channel algorithm to a 16-microphone super-directional beamformer, using both objective measures and human psychophysics, and we showed that listeners received similar amounts of benefit from the proposed algorithm as the super-directional beamformer. Like the auditory system, the proposed algorithm only requires two input channels and does not require *a priori* knowledge of the environment. Our algorithm overcomes specific limitations faced by current state-of-the-art technologies, and provides an alternative, biologically based approached to the cocktail party problem.

## Design and Implementation

### Algorithm

The proposed sound-segregation algorithm uses the neural responses of spatially tuned neurons within a modified model of the barn owl inferior colliculus to segregate sound mixtures [35]. The algorithm contains three modules: peripheral filtering, midbrain model, and stimulus reconstruction (Figure 1). All modules are implemented in MATLAB (Mathworks, Natick, MA). The algorithm is available upon request.

**Figure 1:**
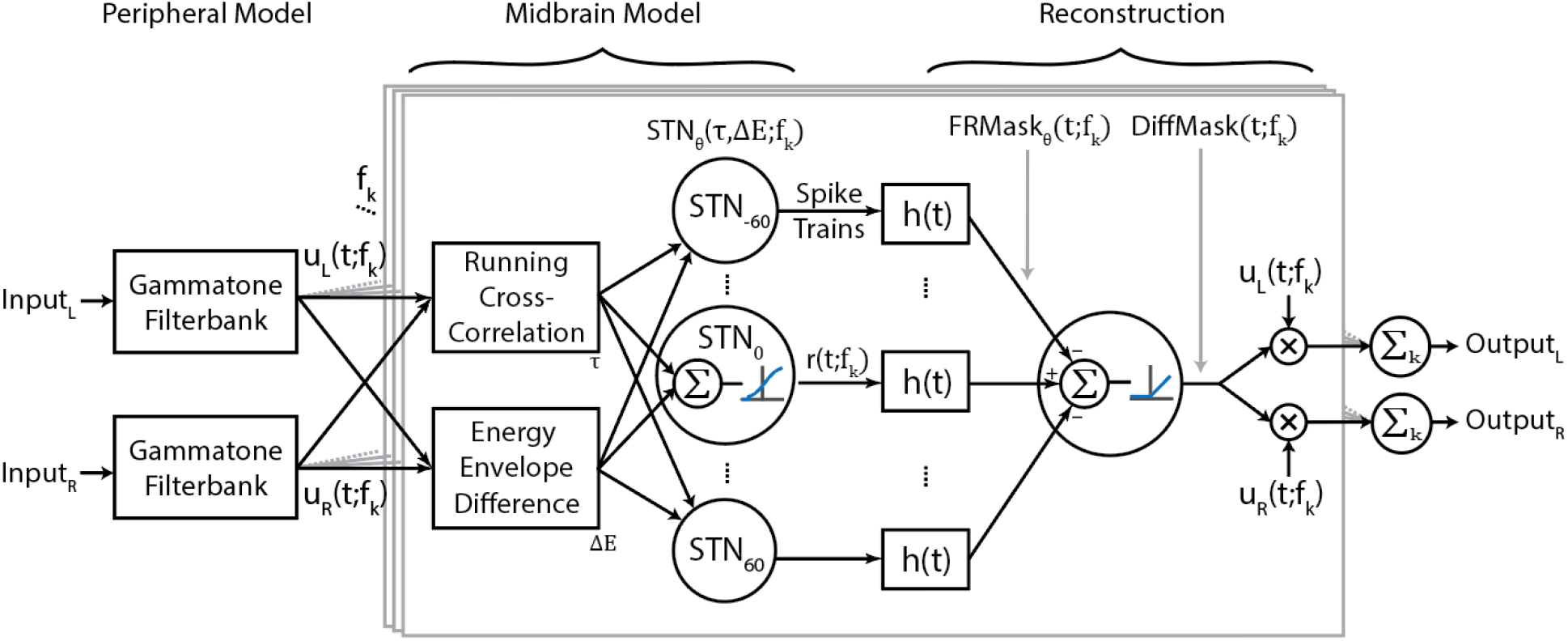
Flow diagram of the proposed algorithm. Central boxes, outlined in grey, show processing for a single-frequency band. The functions u_L_(t; f_k_) and u_L_(t; f_k_) are the narrowband signals of the left and right input channels for each frequency channel, and f_k_ denotes the k^th^ frequency channel. The midbrain model is based on spatially tuned neurons (STNs), where each STN has a “best” ITD and ILD, denoted τ and ΔE, respectively. The best ITD and ILD values of a neuron depend on the direction θ and frequency f_k_ to which the STN is tuned. h(t) represents the reconstruction kernel that converts spike trains to waveforms. The implementation of DiffMask in our analysis involves five sets of STNs, where θ ∈ {0, ±30, ±60}; however, other implementations of the model may involve different sets of θ.

### Peripheral Filtering

Left and right channels of the input audio are filtered with a gammatone equivalent-rectangular-bandwith (ERB) filterbank, implemented using the auditory toolbox in MATLAB [36]. The parameters of ERB spacing were: Q factor = 9.26449 [37], minimum bandwidth = 24.7 Hz, order = 1. The filterbank used here has 64 channels with center frequencies ranging from *f*_1_ = 200*Hz* to *f*_64_ = 20 *kHz*. The filterbank outputs are two sets of 64 channels of narrowband signals, *u*_*L*_*(t; f*_*k*_ ) and *u*_*R*_*(t; f*_*k*_ ), corresponding to the left and right channels, respectively.

### Midbrain Model

The midbrain model represents stimulus-driven sound segregation that occurs in the subcortical regions of the brain. Sound streams from various spatial directions were encoded by populations of *spatially tuned neurons* (STNs), where each neuron is sensitive to one specific direction *θ* in the horizontal plane (*STN*_*θ*_, Figure 1). Within each population, there were 64 neurons tuned to each f_*k*_ of the previous module. Unless otherwise noted, five sets of STNs were constructed in the current study, corresponding to five spatial directions: *θ* ∈ {0°, ±30°, ±60°}. These five directions were chosen to optimize the spatial tuning of the DiffMask reconstruction method (see *Reconstruction*).

Each neuron’s preferred location is determined by its internal *τ* and *ΔE* parameters, corresponding to its best ITD and best ILD, respectively (Figure 1). *ΔE* is frequency dependent, and each neuron’s ΔE was calculated by taking the energy envelope difference [35] of the narrowband Head Related Transfer Functions (HRTFs) of the Knowles Electronic Manikin for Acoustic Research (KEMAR) [38,39] centered around *f*_*k*_. Each neuron’s *τ* was calculated using the Woodworth formulation [40], with the approximation that ITDs are independent of frequency. Preliminary studies found that using frequency-dependent ITD values, calculated as described by the Fischer model or the ones described by Aaronson and Hartmann (2014), provided no benefit in terms of objective measures of algorithm performance (see *Reconstruction*).

The neurons were driven by a combination of two quantities. The first was a short-time running cross correlation (RCC) between the energy-normalized *u*_*L*_(*t*; *f*_*k*_) and *u*_*R*_(*t*; *f*_*k*_), calculated at the neuron’s time-lag T over 5-ms windows. The second was the energy envelope difference (EED) between *u*_*L*_(*t*; *f*_*k*_) and *u*_*R*_(*t*; *f*_*k*_). If the stimulus EED was within the neuron’s specified range of *ΔE*, it was then weighted by the energy envelope of either *u*_*L*_(*t*; *f*_*k*_) or *u*_*R*_(*t*; *f*_*k*_), and multiplied by the RCC. The instantaneous rate of firing of the STN is then a rectified version of this product. Specifically, the RCC-EED combination was transformed via a sigmoidal input-output nonlinearity (i.e., an activation function) to obtain the estimated instantaneous firing rate of the STN. Finally, a Poisson spiking generator was used to generate spike trains for each neuron (*r*_*θ*_ (*t*; *f*_*k*_), Figure 1). For detailed description of the mathematics of these operations and their physiological basis, we refer interested readers to Fischer et al. (2009). Parameters for the input-output nonlinearity were modified to maximize the objective intelligibility of the sounds recovered from the neurons’ corresponding spiking patterns. The output at this stage for each set of STNs is 64 spike trains, corresponding to each frequency channel.

### Stimulus Reconstruction

Each (*r*_*θ*_ (*t*; *f*_*k*_) essentially captured the energies of sound sources originating from *θ*, encoded as neural spikes. The stimulus reconstruction module decoded this information into audible waveforms, using an approach similar to ideal time-frequency mask estimation [42]. The concept of time-frequency masks can be summarized as follows: for a time-frequency representation of an audio mixture (e.g., spectrograms) consisting of a target and interferers, one can evaluate each element (i.e., time-frequency tile) of such a representation and determine whether the energy present is dominated by the target or the masker. If the target sound dominates, a value of unity (1) is assigned to that time-frequency tile, and zero (0) otherwise. This process creates an ideal binary mask. Alternatively, assigning the ratio of energies of the target to total energies in a time-frequency tile yields the ideal ratio mask [43]. One can then estimate the target sound by applying the mask to the sound mixture via element-wise multiplication. This process has been shown to recover the target with high fidelity in various types of noise [42].

The reconstruction methods used here are described in the next two subsections. These methods estimate time-frequency masks based on the spiking patterns of the STNs. Objective measures of intelligibility (STOI) and quality (PESQ) were used to determine optimal parameters for the reconstruction algorithm [44,45].

### Firing Rate Mask

To produce a time-frequency mask from the STNs’ spiking responses, (*r*_*θ*_ (*t*; *f*_*k*_) were convolved with a kernel, *h*(*t*), to calculate a smoothed, firing-rate-like measure. We set the kernel to be an alpha function: 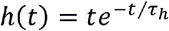, a common function for modeling neural dynamics. The same kernel was convolved with the spike trains of each frequency channel independently. The resulting firing rates of each set of STNs was treated as a non-binary time-frequency mask, which we called “FRMask”:

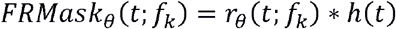

where *θ* denotes the location to which the STNs are tuned. The parameter *τ*_*h*_ was set to 18 ms, and was chosen to maximize the STOI of the final reconstructions. The kernel itself was restricted to a length of 100 ms.

To isolate sounds originating from location *θ*, we applied (i.e., point-multiplied) *FRMask*_*θ*_ *independently* to the left and right channels of the original sound mixture. Then, we summed (without weighting) each frequency channel of the FRMask-filtered signal to obtain an audible, segregated waveform. We designated this result as *Ŝ*_*FR*_.

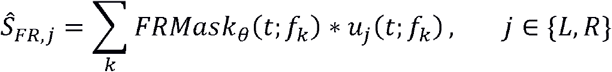

This procedure resulted in a binaural signal, and retained the natural spatial cues of the sound sources.

### Difference Mask

Due to spatial leakage (Figure 2 & Figure 3), the mask-filtered signal obtained using FRMask_0_ inevitably contained information from sounds originating from other spatial locations. To address this overlap in spatial tuning, scaled versions of *FRMask*_±30_ and *FRMask* _±60_ are subtracted from FRMask_0_, followed by half-wave rectification, to produce a *Difference Mask* (“DiffMask”). This operation represents inhibition of the 0° STNs by the ±30° and ±60° STNs:

**Figure 2:**
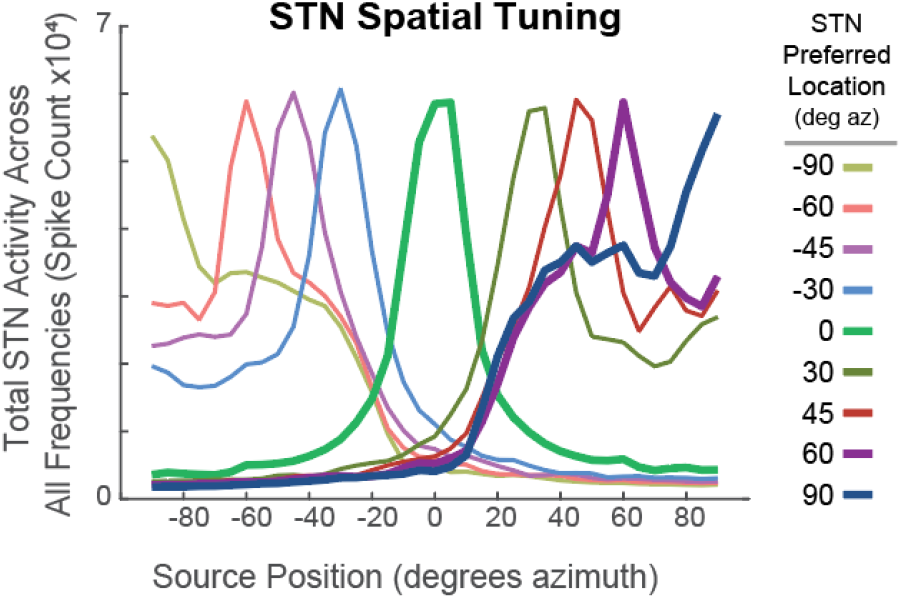
Spatial Tuning Curves of the midbrain model, represented as total neural activity in response to a White Gaussian Noise. Each colored line represents the total response from a set of STNs. Tuning curves of STNs tuned to 0°, 45°, and 90° azimuth are bolded.

**Figure 3:**
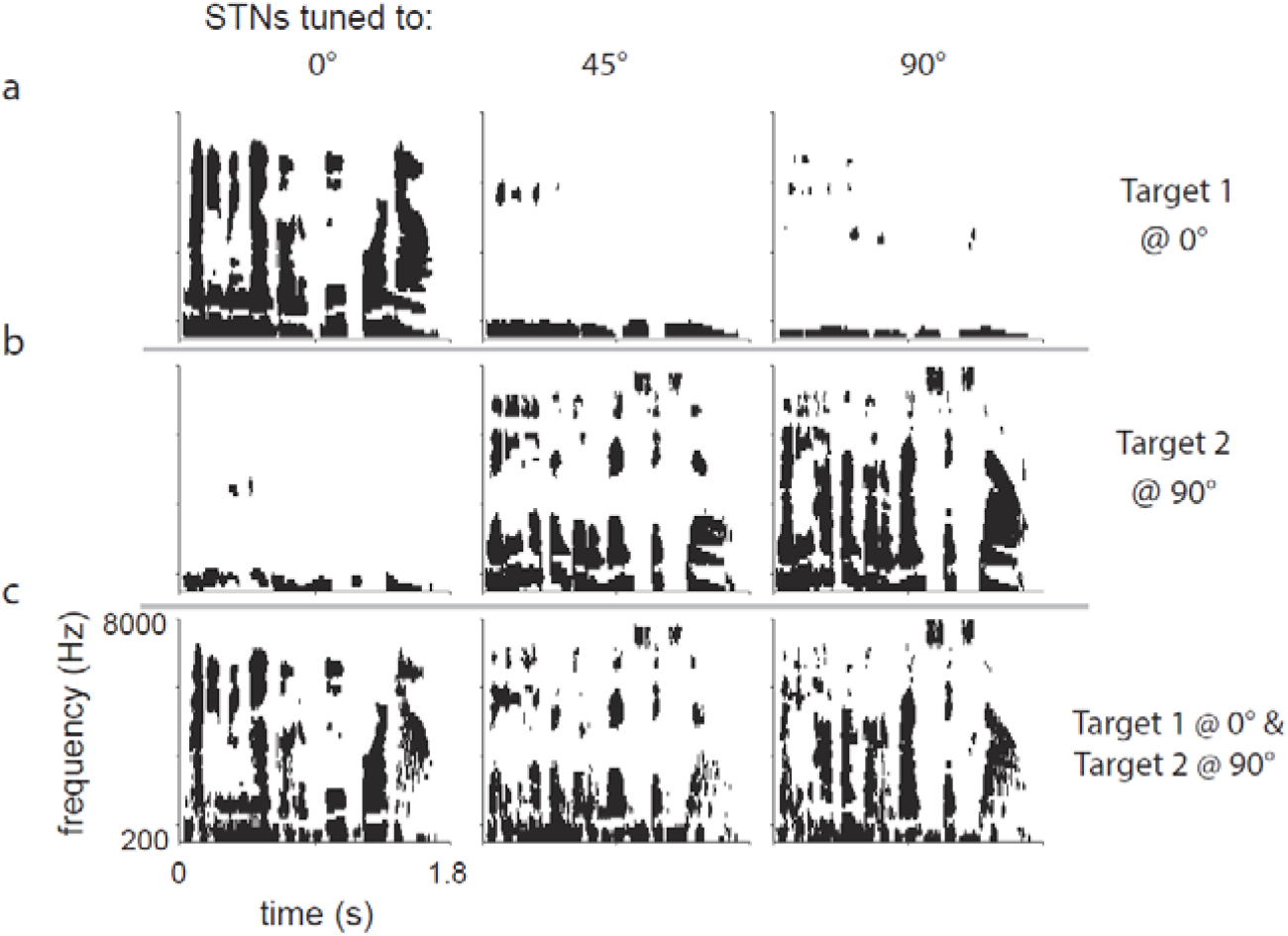
Raster plots of STN responses to (a) a single sentence placed at 0° azimuth, (b) a different sentence placed at 90° azimuth, and (c) both sentences present at their respective locations. Columns represent STNs tuned to the location indicated.

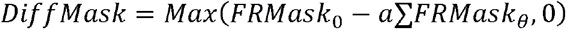

where *θ* ∈ [±30°, ±60°], and was chosen to minimize the spatial leakage of 0° STNs (Figure 4). In the DiffMask operation, all masks are first normalized to have values within [0,1]. The scaling factor a was chosen to be 0.5 to maximize the average STOI of the reconstructed waveforms. The DiffMask is used to reconstruct an audible waveform using the same procedure as the FRMask, resulting in a segregated, binaural waveform which we designated *Ŝ*_*DM*_. In this implementation, the scaling parameter *α* was chosen to be constant across all frequencies and spatial channels to reduce the amount of computational complexity in the algorithm.

**Figure 4:**
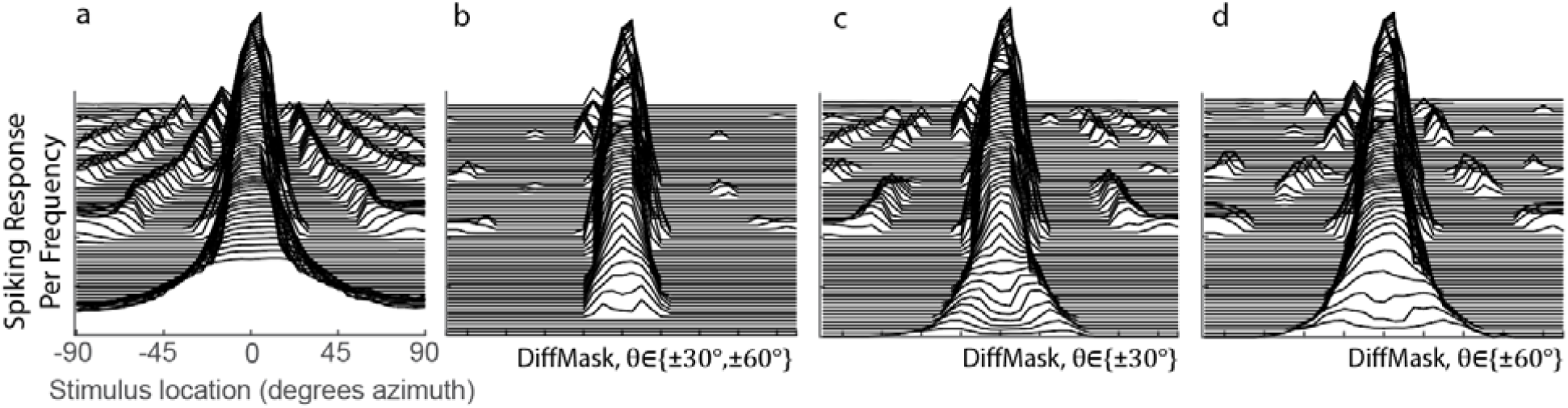
Spatial Tuning of the 0° STNs for before (a) and after (b-d) the DiffMask operation. Each line represents the spatial tuning curve of a single frequency-specific neuron within the set of STNs. STN of the neuron tuned to the lowest frequency is placed on the bottom of the plots. STNs involved in the DiffMask operation are denoted in each subplot.

## Results

### Spatial Tuning Characteristics

Spatial tuning responses of STNs were important predictors of the model’s segregation performance, and helped guide the design of DiffMask. We define “spatial tuning curves” as the spiking activity of STNs as a function of stimulus location. To construct spatial tuning curves, white Gaussian noise was convolved with anechoic KEMAR HRTFs, then presented to the proposed algorithm. Figure 2 shows the responses of STNs combined across frequency channels. Ideally, STNs would only respond to stimuli from one specific direction. However, Figure 2 also shows that all STNs respond to off-target locations. For example, STNs tuned to 0° azimuth (Figure 2, green curve) has a non-zero response to stimuli at ±90° azimuth. We refer to this property as “spatial leakage”, which occurs due to the fact that stimuli from various locations and frequencies share the same ITDs and/or ILDs, making these cues ambiguous indicators of source location [46].

### Spatial Leakage

Leakage across spatial channels limits the performance of the algorithm, especially when multiple sound sources are present. To demonstrate, two randomly selected sentences were presented individually to the model from 0° azimuth (Figure 3a), 90° azimuth (Figure3b), or simultaneously from both locations (Figure 3c). The responses of three set of STNs, tuned to 90°, 45°, and 0°, are shown as spike-rasters. Each row within a raster plot represents the spiking response from the neuron tuned to that particular frequency channel. Due to spatial leakage, all STNs respond to the single sentence placed at 0° or 90° (Figure 3a,b). When both sentences are present, ITDs and ILDs interact to produce complicated STN response patterns (Figure 3c). Spatial leakage limits the ability of STNs to respond to a single talker, since any one spatial channel contains information from other spatial channels. DiffMask was designed to address the issue of spatial leakage by suppressing neural activation by off-target sound streams.

### DiffMask

The DiffMask operation was applied to the spatial tuning curves of 0° STNs to illustrate its sharpening effect on spatial tuning. Figure 4a shows the tuning curves prior to the DiffMask operation. Some neurons within the 0° STNs were activated by stimuli from as far away as 90° (see side peaks). After the DiffMask operation, spiking activity elicited by far-away stimuli were silenced, and side peaks were suppressed considerably (Figure 4b). Using a subset of STNs during the DiffMask operation, such as those tuned to ±30° (Figure 4c) or ±60° (Figure 4d), did not suppress side-peaks as effectively as if both ±30° and ±60° were used.

### Stimulus Reconstruction from Spike Raster Ensembles

So far, we described an algorithm for segregating spatially distributed sound sources within a mixture, and characterized its spatial response properties. We introduced the idea of FRMask and DiffMask for reconstructing neural spike ensembles into audible waveforms. Here, we illustrate and compare these processing methods. One of the 5-dB target-to-masker ratio (TMR), five-talker mixtures described in *Methods* is processed with FRMask and DiffMask. The spectrograms of the resulting reconstructions and their STOIs are shown in Figure 5. Processing the mixture using FRMask provided some segregation benefit over the unprocessed mixture (10% increase in STOI). Additional processing using DiffMask to address spatial leakage provided additional segregation benefit (8% increase in STOI).

**Figure 5:**
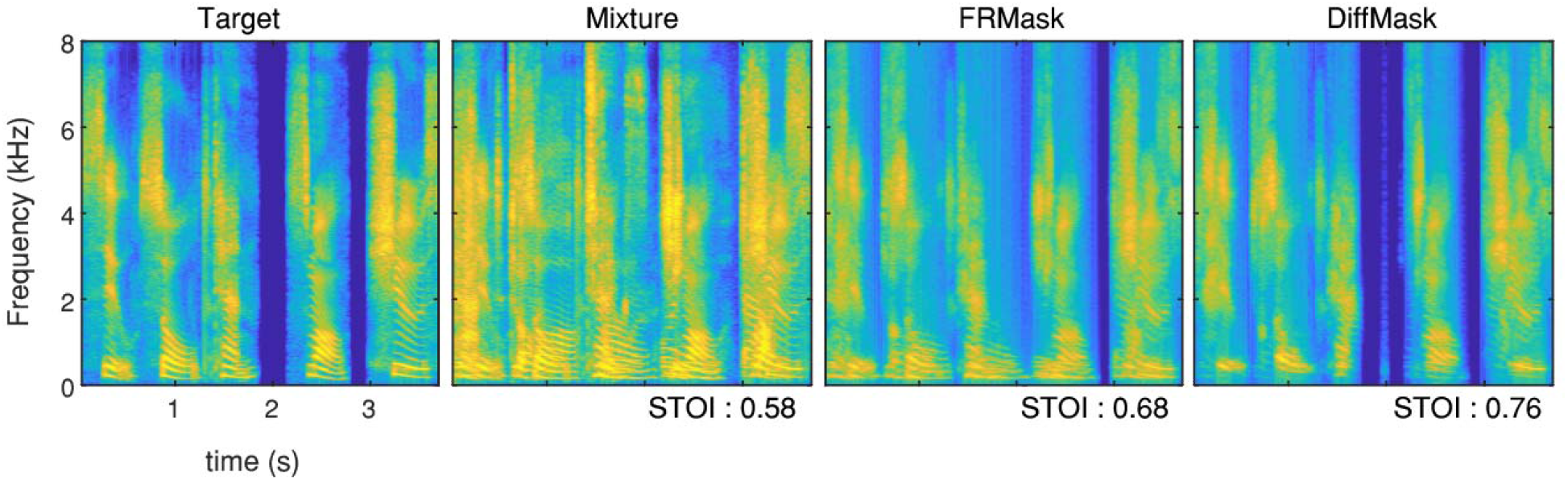
Examples outputs of the various processing methods. A target sentence is embedded in a mixture with four other talkers at 5 dB TMR. Two processing methods are used to recover the target: FRMask and DiffMask. Energies of each example and its corresponding STOI scores are shown.

We also used a different set of stimuli to test the DiffMask in a novel acoustic scene, which were identical as that used by Chou and colleagues to evaluate the PA [34]. The stimuli were 0 dB TMR mixtures of three sentences, where two masker sentences were placed symmetrically around a target sentence at various azimuthal separations. All sentences were spoken by the same talker from the coordinate response measure corpus (Talker 4, arbitrarily chosen), which contained sentences with *different* semantics and temporal patterns than the corpus described in *methods* [47]. Parameters of DiffMask remained fixed, as per the main analysis. Unlike the PA, the DiffMask reconstruction method was able to recover temporal fine structures of the target stimulus (Figure 6a), resulting in a more natural-sounding waveform. Both the PA and the proposed algorithm plateaued in segregation performance when maskers were separated by 20° or more from the target (Figure 6b). However, on average, BOSSA with DiffMask reconstruction outperformed the PA by 12% STOI for this three-stream mixture task.

**Figure 6:**
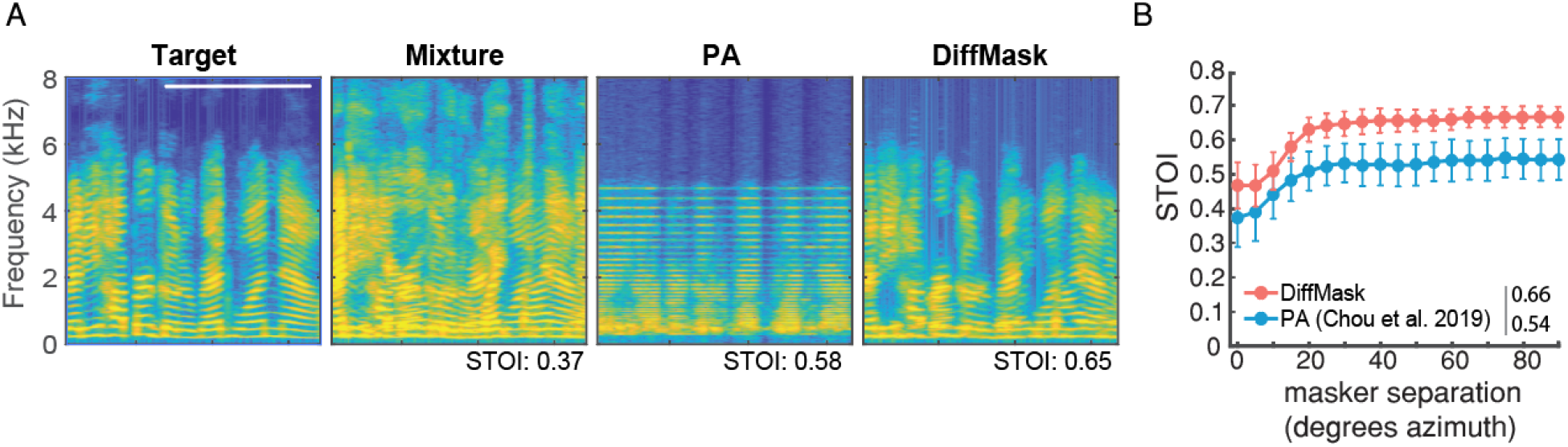
Comparison against the PA. a) Spectrograms of reconstructions using PA and DiffMask on the same three-sentence mixtures. White bar denotes one second. b) Objective intelligibility (STOI) of BOSSA with DiffMask reconstruction versus the PA in segregating 0-dB TMR three-sentence mixtures from the CRM corpus. Average STOI scores for masker separation between 20 and 90° azimuth are shown in the figure legend.

### Psychophysical Results

A psychophysical study (see *Methods*) was conducted to determine the perceptual benefit that BOSSA can provide to listeners. In the study, the two reconstruction methods employed by BOSSA (FRMask and DiffMask) was compared against a 16-microphone super-directional beamformer (BEAMAR, see *methods* for details). Young, normal-hearing listeners were asked to indicate the target words embedded in a 5-sentence mixture on a graphical user interface. Figure 7a shows the results of the task in terms of the percentage of correct responses for each TMR and processing algorithm. Two-way repeated-measures ANOVA found significant interaction between processing method and TMR on performance (F(6,60)=6.97,p<0.001). Post-hoc pairwise comparisons using Tukey’s HSD test found significant differences between performance under the natural condition and under each of the three processing methods for all three TMRs (p<0.001), suggesting that subjects significantly benefitted from listening to processed speech across all TMRs. The relative performance between each algorithm is TMR-dependent. At 5-dB TMR, subjects scored equally high under all three processing conditions. However, at −5- and 0-dB TMR, subjects scored higher with DiffMask than with FRMask, and scored similarly between DiffMask and BEAMAR.

**Figure 7:**
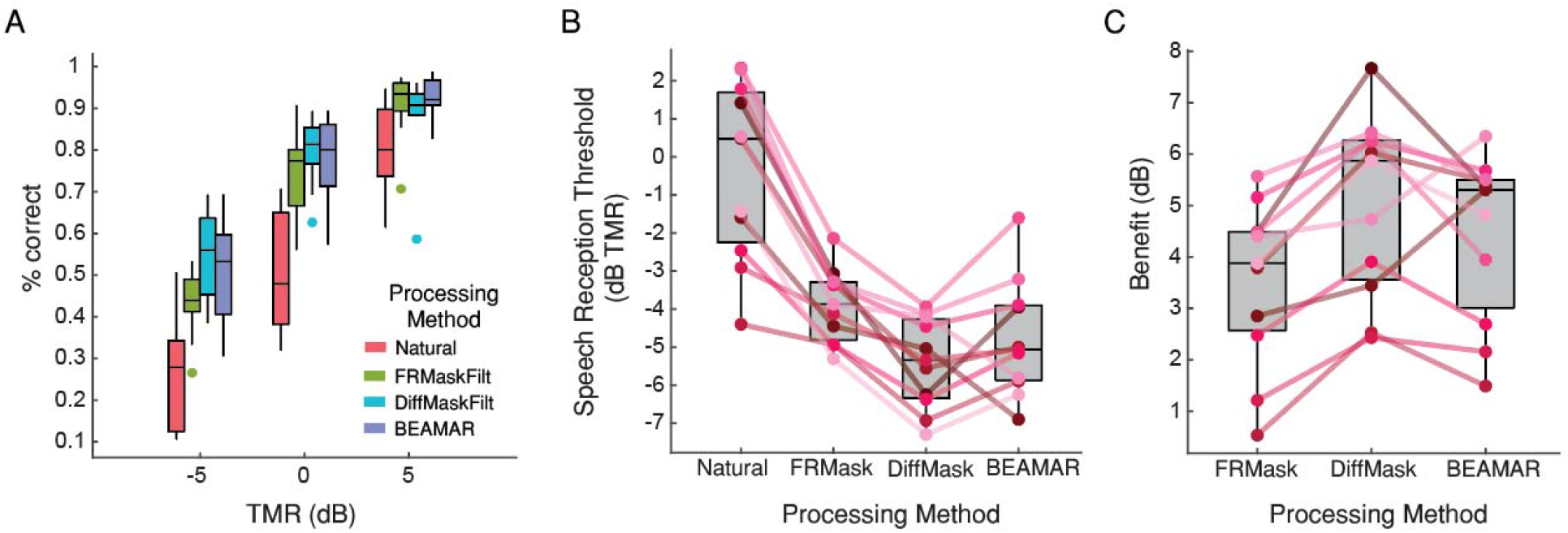
Behavioral evaluation results. a) Average percent correct for each processing condition as a function of TMR. Higher is better. b) Average and individual subject speech reception threshold for each processing method. Lower is better. Solid lines mark the SRT for each participant for each processing condition. c) Average and individual subject perceptual benefit relative to the natural condition. Higher is better.

Figure 7B presents the same results in terms of speech reception thresholds (SRT), defined as the TMR for a listener to correctly respond to 50% of the stimuli, for each processing condition. Each solid circle represents the SRT for a single subject. The perceptual benefit provided by an algorithm is the difference between its SRT and the unprocessed SRT (Figure 7c). ANOVA followed by Tukey’s multiple pairwise comparison showed that all three algorithms provided significant benefit to listeners over the unprocessed condition (p<0.001). Benefits provided by BEAMAR and DiffMask are not significantly different (p=0.66). Two of the eleven listeners two achieved the lowest SRT and gained the most benefit from BEAMAR, while nine out of eleven listeners achieved the lowest SRT and gained the most benefit from DiffMask.

## Discussion

Extensive research has been devoted to developing a solution for the CPP (for review, see Qian et al. 2018), [3] and many approaches benefit from using multiple microphones. For example, the performance of methods using independent component analysis degrades quickly as the number of sources exceeds the number of microphones [48]. In acoustic beamforming, performance of the beamformer can be significantly improved by increasing the number of microphones used [23,24]. Although traditional beamformers produce a single-channel output, which cannot carry binaural information, recent advances in spatial cue preservation strategies were able to overcome this limitation [28,29]. Based on these observations alone, one might predict that in general, algorithms employing microphone arrays would outperform two-microphone algorithms.

On the contrary, by employing deep learning methods such as deep clustering [49] and permutation invariant training [50,51], CASA algorithms have achieved impressive results under controlled environments [52]. Consequently, the domain of CASA as a whole has moved toward using deep learning to perform classification on acoustic features [3,32,52]. However, deep learning approaches have several limitations: 1. Though the features used may be biologically relevant, the classification algorithm itself has no biological interpretation; 2. They require supervised training with large datasets; 3. Due to their dependence on supervised training, the generalization capability of DNNs remain unclear. The final limitation may be a very important factor in the adoption of DNNs in hearing assistive devices, which are often challenged by a variety of novel acoustic scenes.

Rather than using deep learning, this study presents a binaural CASA algorithm that uses a model of the auditory system to perform classification. There are several advantages to this. First, the proposed algorithm does not require training and is likely to generalize well to novel datasets. Figure 6 shows that the proposed algorithm performs well in a novel acoustic scene, which was composed of a speech corpus with different speakers, semantics, temporal patters, number of talkers, and talker positions than the one described in *Methods*. Next, an important and unique advantage of the proposed algorithm is its biological interpretability. The role of every single neuron involved in the proposed algorithm can be explained by their spatial tuning characteristics (Figure 2). Unlike DNNs, the design of BOSSA can be easily adapted to incorporate novel mechanisms observed in neurophysiology.

When compared to the unprocessed condition, the proposed algorithm provided a significant amount of benefit to normal-hearing listeners in a difficult listening task (Figure 7). While BEAMAR and the proposed algorithm provided a similar amount of benefit, the majority of listeners benefitted the most from the proposed algorithm. This is an extremely promising result, considering that BEAMAR uses 14 more microphones than the proposed algorithm.

### Relationship with Biology

One advantage of the proposed algorithm is that is can be related to observed biological phenomena. Interestingly, the spatial tuning curves of the STNs in our model, combined across frequency channel, (Figure 2a, bolded lines) resemble the spatial tuning curves of “center-preferred” and “contra-preferred” neurons found in the auditory cortex of mice [53,54], cats [55], rhesus monkeys [56], and humans [57]. The similarities between the experimentally observed tuning curves and the modeled center-preferred and contra-preferred tuning curves suggest the emergence of similar mechanisms behind spatial tuning properties across multiple species.

In our model, sharpening of spatial tuning curves resulted in a significant increase in segregation performance. The DiffMask operation responsible for this can be described by a lateral inhibition network. Such a network is commonly found in biological systems, and also play a role in sharpening spatial and temporal input patterns [58].

### Performance versus Beamformers

Fixed beamforming algorithms, such as BEAMAR, attenuate off-target sounds by combining the delayed and weighted signals recorded at each microphone. This operation takes advantage of the differences in arrival times of the target signal at each receiver. In contrast, BOSSA makes use of both timing and level differences of the binaural signal to perform segregation. The additional use of ILDs may partially explain why a binaural algorithm can achieve performance parity with a 16-microphone algorithm. This hypothesis is consistent with physiological studies which show the importance of ILD cues for sound localization in humans and other mammals [59,60].

### Time-Frequency Mask Estimation from Spike Raster Ensembles

The firing rates (FR) calculated from the midbrain model are used to calculate spike trains. These spike trains are then used to calculate FRMask, a firing rate-like measure. Comparing FR with FRMask, we find that FRMask is akin to a smoothed version of FR. In theory, FRMask and DiffMask can be directly derived from FR, making the calculation of spike rasters seemingly redundant. However, since the midbrain model can be used as a front-end to spiking network models, the calculation of spikes is necessary for those applications [34]. In this study, we demonstrated that the proposed method of mask-estimation yields highly intelligible reconstructions from spiking neurons. This method enables one to analyze each node within spiking networks and obtain intelligible reconstructions at each node.

Spiking neural networks traditionally do not have applications in audio processing due to the lack of a method that produces intelligible, high quality reconstructions. The “optimal prior” method of reconstruction is often used to obtain reconstructions from physiologically recoded neural responses [61– 64], but produces single-audio-channel reconstructions of poor quality and intelligibility [34]. The optimal prior method computes a linear filter between a training stimulus and the response of neuron ensembles, and filter needs to be re-trained if the underlying network changes. In contrast, the mask-based reconstruction method used in this study estimates time-frequency masks from spike trains. It is able to obtain reconstructions with much higher intelligibility and preserves spatial cues, all without the need for training (Figure 6). These properties enable rapid development of spiking neural network models for audio-related applications.

### Limitations and Future Work

While the formulation of DiffMask can sharpen the spatial tuning of the STNs, neurons tuned to frequencies below 300 Hz were completely silenced (Figure 4b). Additionally, some side peaks still persist even after the DiffMask operation, implying that spatial leakage was not fully addressed (Figure 4b). Different formulations to the DiffMask should be explored to address these shortcomings. We have avoided using deep-learning approaches in this study in favor of biological interpretability, but such approaches may help guide the optimization of DiffMask, and should be considered in the future. Next, the algorithm was designed to process each frequency channel independently. This design choice reduces both the complexity of the algorithm and its computation time. However, cross-frequency connections certainly take place in the auditory system, as spectral grouping of perceptually relevant sounds has been observed in many animals, and may explain the missing fundamental phenomenon (see discussion by Medvedev, Chiao, and Kanwal 2002). To further improve the algorithm performance, interactions across frequency channels should be explored. Finally, animals have been observed to resolve binaural cue ambiguity by having neurons preferentially tune to more reliable spatial cues [66]. Such physiological approaches to overcome spatial leakage should also be considered. Incorporating deep-learning based optimization methods may help identify these “reliable” cues.

## Methods

### Psychophysics Evaluation

A psychophysical study was conducted in order to determine the subjective benefit provided by the algorithm. The performance of FRMask and DiffMask was compared against a 16-microphone super-directional beamformer, called BEAMAR [27,28].

BEAMAR attenuates off-center sounds by combining the weighted output of 16 omni-directional microphones into a single channel, using an optimal-directivity algorithm [27,67]. The microphones were arranged in columnar groups across the top of the head [68]. Since the presence of natural spatial cues has been shown to provide a large perceptual benefit to listeners (e.g., through spatial release from masking Litovsky, 2012), BEAMAR does not process frequencies below 1 kHz to retain natural spatial cues in that frequency region. The combination of beamforming at high frequencies and natural binaural signals at low frequencies has been shown to provide a significant benefit to both normal-hearing and hearing-impaired listeners attend to a target speech sentence in a multi-talker mixture [28].

### Stimuli

Stimuli used in the objective and behavioral evaluations were designed to prevent listeners from using monaural cues when solving the task, and thus focused the investigation on spatially based segregation. Five-word sentences were constructed from a corpus of monosyllabic words [69], with the form [name-verb-number-adjective-noun] (e.g. “Sue found three red hats”). The corpus contains eight words in each of the five categories. Each word in a sentence was spoken by a different female talker, randomly chosen from a set of eight female talkers, without repetition. During each trial, a target sentence was mixed with four masker sentences, all constructed in the same manner. Repetition of the same talker within a word category was allowed. The randomization of talkers within a sentence was designed to prevent subjects from using the timbre of the talkers as a segregation cue. Words from the target and masker sentences were time-aligned, so that the words from each category shared the same onset. This was designed to prevent the listeners from using temporal cues to segregate the mixture.

The five sentences were simulated to originate from five spatial locations: 0°, ±30°, and ±60° azimuth, by convolving each sentence with anechoic KEMAR HRTFs [38]. The target sentence was always located at 0° azimuth. Each masker was presented at 55 dB sound pressure level (SPL). The level of the target was varied to achieve target-to-masker ratios (TMRs) of −5, 0, and 5 dB.

### Experiment Conditions

Sentences from the five-talker stimuli were processed using one of three methods: BEAMAR, FRMask, and DiffMask. An unprocessed condition, “KEMAR”, was used as a control. In KEMAR, sentence mixtures with natural ITD and ILD cues as derived from KEMAR were presented to the participants. BEAMAR was the 16-microphone acoustic beamforming algorithm described previously. The remaining two processing conditions corresponded to the two reconstruction methods described under *Stimulus Reconstruction*: FRMask and DiffMask.

### Participants

Participants in this study were eleven young normal-hearing listeners, ages 18-32. All listeners had symmetrical audiogram measurements between 0.25 – 8 kHz with hearing thresholds within 20 dB HL. Participants were paid for their participation and gave informed consent. All procedures were approved by the Boston University Institutional Review Board (protocol 1301E).

### Procedures

Three blocks were presented for each of the four conditions, with each block containing five trials of each of the three TMRs (15 total trials per block). This resulted in 15 trials per TMR for each of the four processing conditions, and a total of 180 trials across all conditions. The order of presentation of TMRs within a block, and the order of blocks for each participant, were chosen at random. The experiment took approximately one hour to complete.

Stimuli were controlled in MATLAB and presented via a real time processor and headphone driver (RP2.1 & HB7, Tucker Davis Technologies) through a pair of headphones (Sennheiser HD265 Linear). The sound system was calibrated at the headphones with a sound meter (type 2250; Brüel & Kjær, Nærum, Denmark). Participants were seated in a double-walled sound-treated booth. A computer monitor inside the booth displayed a graphical user interface containing a grid of 40 words. For each trial, participants were presented a sentence mixture and were instructed to listen for the target sentence located directly in front. Participants responded with a mouse by choosing one word from each column on the grid.

### Analysis

Each participant’s performance was evaluated by calculating the percentage of correctly answered keywords across all trials. Psychometric functions were generated for each listener in each condition by plotting the percent correct as a function of TMR, and fitting a logistic function to those data. Speech reception thresholds (SRTs), which are the TMRs at which participants scored 50% correct, were extracted from each function using the psignifit toolbox [70]. Differences in SRTs between the natural condition (“KEMAR”) and each of the processing conditions is the “benefit” provided by that processing method. Statistical analysis was done in Python using the statsmodels package [71]. Percent-correct responses were evaluated using two-way repeated-measures ANOVA, followed by Tukey’s HSD test for pairwise comparison. Group means of SRTs and processing benefit were evaluated using one-way repeated-measures ANOVA.

## Acknowledgements

The authors would like to thank Matthew Goupell for reviewing the manuscript.

## Author Contributions

KC designed the algorithm. KC and VB designed the experiment. KC conducted the experiment and analyzed the data. KC wrote the paper. KS and SC reviewed the manuscript and supervised the study.

